# Neurovascular mitochondrial susceptibility impacts blood-brain barrier function and behavior

**DOI:** 10.1101/2024.02.15.580544

**Authors:** A. M. Crockett, H. Kebir, M. C. Vélez Colόn, D. M. Iascone, B. Cielieski, A. Rossano, A. Sehgal, S. A. Anderson, J. I. Alvarez

**Affiliations:** Department of Pathobiology, School of Veterinary Medicine, University of Pennsylvania; Department of Child and Adolescent Psychiatry, Children’s Hospital of Philadelphia; Chronobiology and Sleep Institute, Perelman School of Medicine, University of Pennsylvania; Neurobehavioral Core, Perelman School of Medicine, University of Pennsylvania; The Howard Hughes Medical Institute, Perelman School of Medicine, University of Pennsylvania; Department of Psychiatry, Perelman School of Medicine, University of Pennsylvania

**Keywords:** Blood-brain barrier, 22q11.2 deletion syndrome, mitochondria, neurovascular, and behavior

## Abstract

Maintenance of blood-brain barrier (BBB) integrity is critical to optimal brain function, and its impairment has been linked to multiple neurological disorders. A notable feature of the BBB is its elevated mitochondrial content compared to peripheral endothelial cells, although the functional implications of this phenomenon remain unknown. Here we studied BBB mitochondrial function in the context of the 22q11.2 deletion syndrome (22qDS), a condition associated with a highly increased risk for neuropsychiatric disease. As the 22q11.2 deletion includes 6 mitochondrial genes, and because we have previously identified BBB impairment in 22qDS, we addressed the hypothesis that mitochondrial deficits contribute to BBB dysfunction and impact behavior in this condition. We report mitochondrial impairment in human induced pluripotent stem cell (iPSC)-derived BBB endothelial cells from 22qDS patients, and in BBB endothelial cells from a mouse model of 22qDS. Remarkably, treatment to improve mitochondrial function attenuates mitochondrial deficits and enhances BBB function in both the iPSC and mouse 22qDS models. This treatment also corrected social memory in the mouse model, a deficit previously associated with BBB dysfunction. As BBB integrity correlated with social memory performance, together our findings suggest that mitochondrial dysfunction in the BBB influences barrier integrity and behavior in 22qDS.

## Introduction

Brain homeostasis requires the proper functioning of the blood-brain barrier (BBB), composed of central nervous system (CNS) endothelial cells (ECs). BBB-ECs exhibit specialized properties that function to segregate the CNS parenchyma from the periphery (*1, 2*). One of these unique features is the expression of an elaborate junctional network that underlies the barrier properties of the BBB. Proper formation of these tight junction complexes is integral to maintain the delicate balance of ionic concentration in the interstitial fluid and to sequester inflammatory mediators and toxins capable of ultimately impacting neuronal health and activity (*3*). Deficits in BBB function are believed to contribute to conditions ranging from neurodevelopmental disorders, such as autism and schizophrenia (SZ), to neurodegenerative conditions including Alzheimer’s and Parkinson’s diseases (*4–8*). Indeed, BBB impairment has been directly linked to neuropsychiatric-like symptoms, and breakdowns in barrier integrity are associated with detrimental neuroimmune and neurodegenerative consequences (*9–13*). Thus, the BBB represents a broad therapeutic target for neurological diseases throughout the lifespan and understanding how the BBB maintains its barrier integrity is integral to advancing treatment options in clinical populations.

While the underlying causes of BBB deficits in neurological conditions are unknown, the role of mitochondrial dysfunction and oxidative stress in the diseased brain has become an area of focus (*14, 15*). Mitochondrial dysfunction has been associated with neurodevelopmental and neurodegenerative disorders and is believed to contribute to their disease course (*15–18*). Yet, little is known about the role of the mitochondria in the BBB. Indeed, ECs generally are considered to rely primarily on glycolysis for their metabolic needs, and as such mitochondrial contributions to BBB function have been underexplored (*19*). However, BBB-ECs present with relatively higher mitochondrial content, suggesting that these cells harbor a specialized metabolic need that relies more on oxidative respiration compared to related cell types (*20*). In support of this theory, multiple studies have identified that genetic or pharmacological interference with mitochondrial respiration impairs BBB integrity (*21–23*). However, it remains unclear whether mitochondrial function in the BBB contributes to disease, and whether this system can be therapeutically targeted to preserve or enhance barrier function.

To address these questions, we have studied the BBB in the 22q11.2 deletion syndrome (22qDS, also known as DiGeorge syndrome and velocardiofacial syndrome). This condition is due to a 3-Mb hemizygous deletion of 42 protein-coding genes on chromosome 22 (*24*), and confers an increased risk for both neurodevelopmental conditions, including autism and attention deficit hyperactivity disorder, as well as neurodegenerative diseases, primarily Parkinson’s disease (*25, 26*). As this population is at a 25-fold increased risk for psychosis, the 22q11.2 deletion represents one of the strongest monogenic risk alleles for SZ (*27, 28*). Notably, each of these conditions have been linked to BBB dysfunction alongside mitochondrial impairment (*4–6, 16, 29, 30*). Based on the inclusion of 6 mitochondrial genes in the deleted region, mitochondrial function in 22qDS has emerged as a major focus of the field (*31, 32*). However, the status of the mitochondria in BBB-ECs in the context of 22qDS is unknown. As we have previously found that the BBB is impaired in 22qDS (*33*), we hypothesized that mitochondrial susceptibility contributes to BBB dysfunction and disease in 22qDS.

Here, we have employed human induced pluripotent stem cell (iPSC)-derived BBB-ECs (iBBBs) from 22qDS patients and age- and sex-matched healthy controls (HC) to study mitochondrial function in the BBB. We further address these questions in a mouse model of 22qDS (22qMc), which harbors a homologous hemizygous deletion (*34, 35*). We report profound deficits in mitochondrial function in 22qDS iBBBs. We identify bezafibrate, a peroxisome proliferator-activated receptor (PPAR) agonist (*36*), as a pharmaceutical intervention that improves mitochondrial respiration, which was associated with enhanced barrier function of the BBB *in vitro*. Notably, we found that bezafibrate improved BBB function in 22qMc and restored performance in a behavioral task associated with neuropsychiatric disorders.

## Results

### Mitochondrial function is impaired in the 22qDS iBBB

The 22q11.2 deleted region includes at least 6 genes that encode for mitochondrial-localizing proteins (*37*), and mitochondrial deficits have been reported in various neuronal cell types in murine and human models of 22qDS (*14, 32, 38, 39*). As BBB-ECs harbor a higher mitochondrial content compared to peripheral ECs (*20*), we hypothesized that these cells may be sensitive to mitochondrial dysfunction in the context of 22qDS. To address whether the 22qDS BBB exhibits mitochondrial dysfunction, we studied mitochondrial respiration in iPSC-derived BBB cells from individuals with 22qDS and SZ along with our iPSC lines from undeleted, neurotypical controls. We utilized Resipher, a novel approach for the repeated measurement of oxygen consumption by cultured cells and observed a significant impairment in the oxygen consumption rate (OCR) in 22qDS iBBBs compared to control iBBBs (Figure 1A-B), indicating deficits in mitochondrial respiration. To further probe the status of the mitochondria in the 22qDS BBB, we performed the Seahorse mitochondrial stress test (Figure 1C) and found significantly worsened spare capacity and maximal respiration in 22qDS iBBBs, with trends towards decreased basal respiration and ATP-linked respiration (Figure 1D). 22qDS iBBBs did not upregulate glycolysis to compensate for deficits in respiration, and indeed these cells also appear to have significant impairment in their extracellular acidification rate (ECAR, Figure 1E). Together, our measures indicate a substantially different energetic landscape in 22qDS iBBBs relative to controls (Figure 1F). Based on these findings, we anticipated decreased ATP production, and indeed we observed a significant deficit in the amount of ATP present in 22qDS iBBBs relative to controls (Figure 1G). As these mitochondrial deficits could be due to decreased mitochondrial number in 22qDS iBBBs, we quantified mitochondrial content by assessing mitochondrial DNA copy number (Figure 1H) and observed no significant difference. Together, our findings indicate that mitochondrial respiration is impaired in the 22qDS iBBB resulting in deficient ATP production, and this does not appear to be due to reduced mitochondrial mass in the cells.

**Figure 1.**
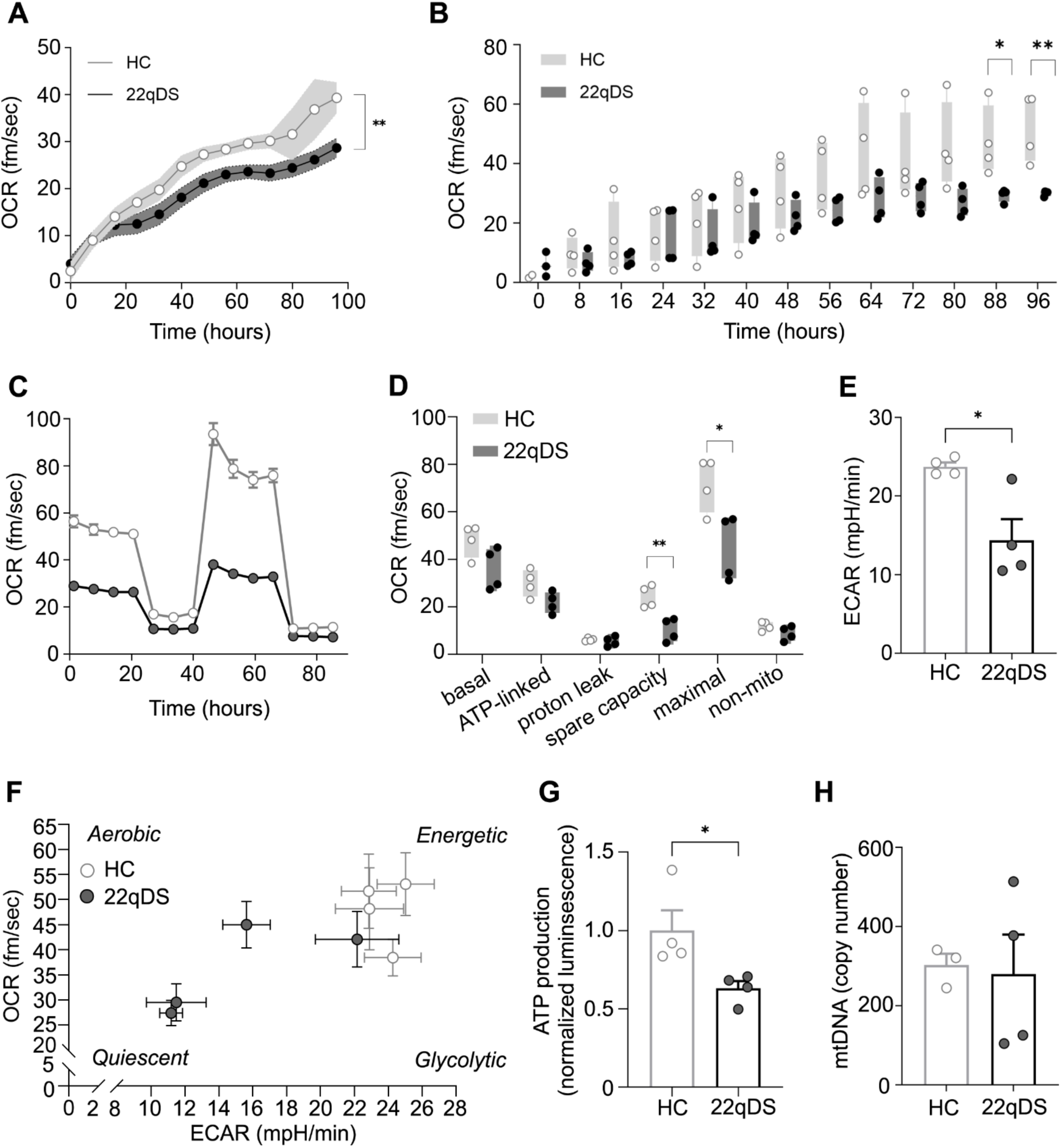
Human 22qDS iPSC-derived BBB cells exhibit mitochondrial deficits. (**A**) Representative time course of real time oximetry monitoring of a one 22qDS iBBB line and the matched control iBBB line obtained using Resipher (*n* = 5-6 replicates per line, two-way ANOVA). Error bars ± SD. (**B**) Quantification of average oxygen consumption rate for each 22qDS and matched control iBBB line over the course of 96 hours (*n* = 4 lines per genotype, 3-6 replicates per line, unpaired t test). (**C**) Representative plot of oxygen consumption rate by one 22qDS iBBB line and the matched control iBBB line during the Seahorse XFe96 Analyzer mitochondrial stress test (*n* = 7-8 replicates per line). (**D**) Quantification of OXPHOS parameters during the Seahorse XFe96 Analyzer mitochondrial stress test for each 22qDS and matched control iBBB line (*n* = 4 lines per genotype, 7-8 replicates per line, unpaired t test). (**E**) ECAR quantification to assess glycolysis by the Seahorse XFe96 Analyzer (*n* = 4 lines per genotype, 7-8 replicates per line, unpaired t test). (**F**) Energy map relating respiration and glycolysis for each iBBB line. Error bars ± SD. (**G**) Quantification of intracellular ATP content (*n* = 4 lines per genotype, 3 replicates per line, unpaired t test). (**H**) Quantification of mitochondrial DNA copy number. Error bars ± SEM unless otherwise indicated, * p < 0.05, ** p < 0.01, *** p < 0.001.

### Targeting mitochondria improves barrier function in 22qDS iBBBs

Interfering with mitochondrial respiration can cause deficits in the barrier integrity of the BBB (*21–23*), but it is unknown whether improving mitochondrial phenotype can restore BBB function. As we have previously reported that the BBB is impaired in 22qDS (*33*), we hypothesized that mitochondrial dysfunction could contribute to the deficit in barrier integrity in this condition. To address this, we identified bezafibrate, a PPAR agonist that was recently reported to attenuate mitochondrial dysfunction in the context of 22qDS (*39*), as a candidate to improve mitochondrial phenotype. We observed that treatment with bezafibrate enhanced mitochondrial function in the Seahorse mitochondrial stress test (Figure 2A-B) without affecting glycolysis (Supplementary Figure 1A). Specifically, bezafibrate treatment significantly improved basal respiration and ATP-linked respiration, suggesting a potential impact of bezafibrate on cell energetics. Bezafibrate also significantly improved maximal respiration and spare capacity, indicating that treatment improved the respiratory range of the 22qDS iBBB. As bezafibrate is a known activator of peroxisome proliferator-activated receptor gamma coactivator (PGC)-1α, which drives mitochondrial biogenesis (*40, 41*), we addressed the hypothesis that bezafibrate functions to increase mitochondrial respiration by enhancing mitochondrial content. To this end, we assessed mitochondrial mass and quantified mitochondrial DNA content. We found a non-significant decrease in Mitotracker fluorescent intensity and a significant decrease in mitochondrial DNA content (Supplementary Figure 1B-C), together indicating that bezafibrate does not improve respiration simply by increasing the amount of mitochondrial mass.

**Figure 2.**
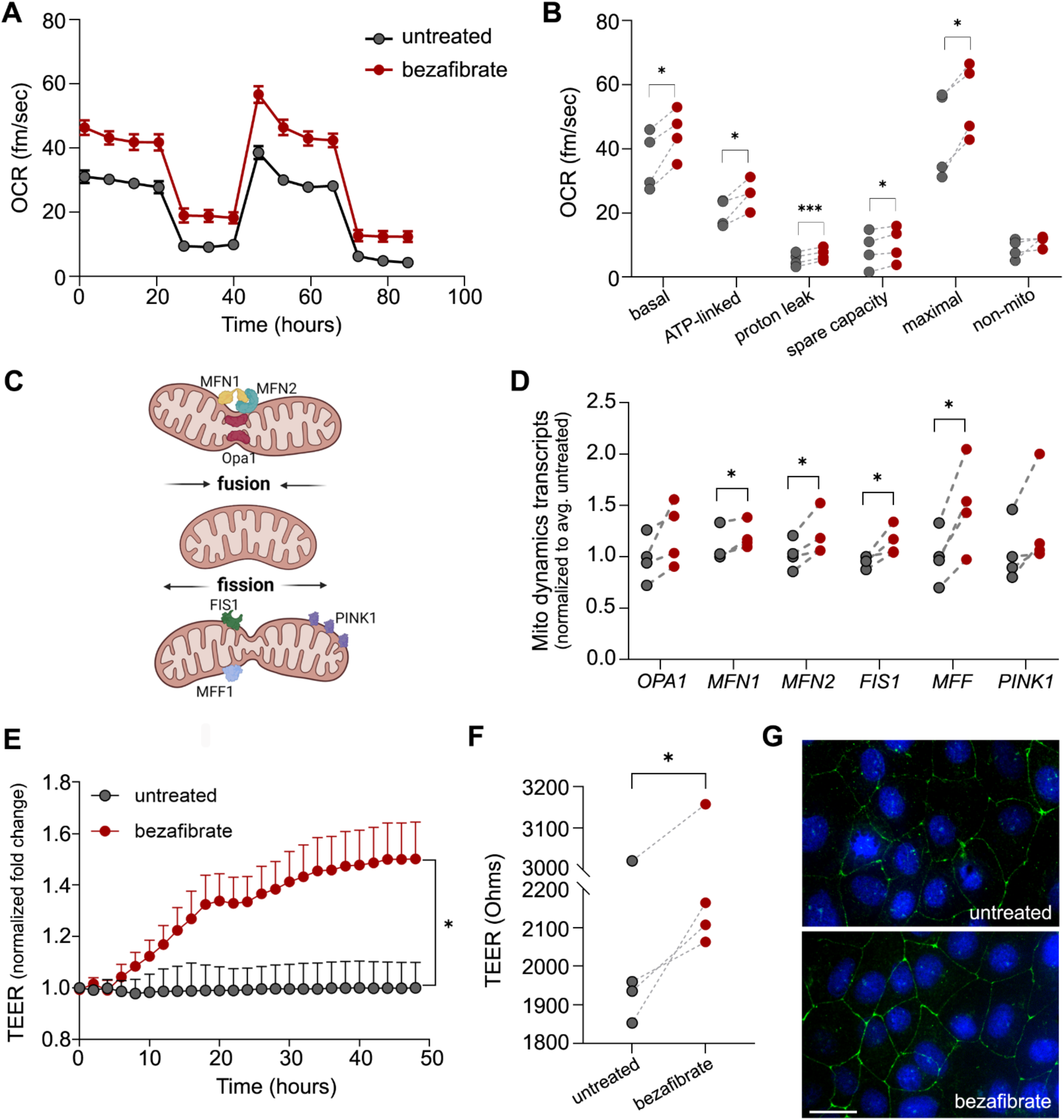
Bezafibrate treatment improves mitochondrial and barrier integrity phenotypes in 22qDS iBBB. (**A**) Representative plot of oxygen consumption rate by a bezafibrate-treated and untreated 22qDS iBBB line during the Seahorse XFe96 Analyzer mitochondrial stress test (*n* = 4 22qDS lines, 7 replicates per condition). (**B**) Quantification of OXPHOS parameters for each 22qDS line comparing bezafibrate-treated and untreated conditions (*n* = 4 22qDS lines, 7 replicates per condition, paired t test). (**C**) Schematic is describing key regulators of mitochondrial dynamics. (**D**) Quantification of gene transcripts for fusion and fission (*n* = 4 22qDS lines, paired t test). (**E**) Representative time course of TEER fold change following bezafibrate treatment in a 22qDS iBBB line, normalized to untreated (*n* = 6-11 replicates per condition, two-way ANOVA). (**F**) Quantification of TEER at 36 hours following bezafibrate treatment (*n* = 4 22qDS lines, 3-4 replicates per condition – 2 experiments per line, paired t test). (**G**) Immunofluorescence of claudin-5 (green) in 22qDS in untreated (top) compared to bezafibrate treatment (bottom). Error bars ± SEM. Scale bar: 10 μm.

A decrease in mitochondrial content suggests an effect on mitochondrial dynamics, the rate of fusion and fission. As altering fusion can impact respiration, we assessed transcripts associated with mitochondrial dynamics (Figure 2C). We found that bezafibrate significantly increased the expression of transcripts associated with mitochondrial fusion, including mitofusin 1 (*MFN1*), and mitofusin 2 (*MFN2*), alongside a non-significant increase in the fusion transcript optic atrophy gene 1 (*OPA1)* (Figure 2D) (*42*). Furthermore, we also identified a significant increase in transcripts associated with mitochondrial fission, including mitochondrial fission protein 1 (*FIS1*) and mitochondrial fission factor (*MFF*) (Figure 2D) (*43, 44*). Together, our findings suggest that bezafibrate acts to improve mitochondrial respiration by affecting the balance of fusion and fission.

To mechanistically link mitochondrial and barrier phenotypes in the 22qDS iBBB, we next determined the impact of bezafibrate treatment on barrier function. Through the transendothelial electrical resistance assay (TEER), we found that treatment with bezafibrate significantly enhanced resistance of the monolayer in the 22qDS iBBB relative to untreated (Figure 2E-F), indicating improved barrier function. As we had previously found that claudin-5 organization is affected in the 22qDS iBBB (*33*), we assessed whether bezafibrate treatment impacted claudin-5 distribution. Through immunofluorescent staining, we observed improvement in claudin-5 structure upon treatment with bezafibrate, relative to the untreated condition (Figure 2G). Together, these findings suggest that improving mitochondrial respiration enhances barrier function in the 22qDS BBB.

22qDS has been previously associated with elevated reactive oxygen species (ROS) accumulation (*14*). Because bezafibrate can have antioxidant effects (*45*), and elevated ROS can impact barrier integrity (*46, 47*), we assessed ROS levels in the 22qDS iBBB. While we found elevated ROS in 22qDS iBBBs (Supplementary Figure 2A-B), we found no effect of bezafibrate on either cellular or mitochondrial ROS (Supplementary Figure 2C-D), suggesting that bezafibrate does not improve barrier integrity by attenuating oxidative stress. To further confirm that the impaired barrier phenotype in the 22qDS iBBB is unrelated to enhanced ROS accumulation, we treated cells with varying concentrations of N-acetyl cysteine, a ROS scavenger (*48*). No effect of this antioxidant treatment on barrier function was detected (Supplementary Figure 2E), supporting the conclusion that barrier integrity deficiency in the 22qDS iBBB is not caused by elevated ROS.

### Bezafibrate treatment improves the mitochondrial and barrier phenotypes of the 22qMc BBB

We next tested our hypothesis that mitochondria are impaired in the 22qDS BBB in 22qMc, the murine model harboring a homologous hemizygous deletion (*34*). We first assessed mitochondrial mass in 22qMc compared to WT, and as *in vitro* we observed no change in mitochondrial content by Mitotracker (Figure 3A). To address mitochondrial phenotype *in vivo*, we used transmission electron microscopy (EM) to study mitochondrial ultrastructure in BBB-ECs. As mitochondrial stress can drive changes in morphology, causing mitochondria to become swollen and rounder, we assessed mitochondrial indices related to their shape or structure. We observed no evidence of mitochondrial dysmorphia, as measured by mitochondrial length and mitochondrial area (Figure 3B-D). However, we noticed that in the absence of overt changes in mitochondrial morphology, the mitochondria in 22qMc BBB-ECs looked unhealthy, as they appeared paler than their WT counterparts. We identified that the intramitochondrial density is significantly decreased in the BBB-ECs from 22qMc relative to WT, suggesting that mitochondria are dysfunctional in the 22qMc BBB (Figure 3B, E). To determine if bezafibrate improves mitochondrial phenotype *in vivo*, we treated mice with 0.5% bezafibrate-enriched chow for 4 weeks and characterized the mitochondria in BBB-ECs isolated from the 22qMc brain. While we found no effect of treatment on mitochondrial content in the 22qMc BBB (Figure 4A), we did observe an impact on intramitochondrial density (Figure 4B-C) by EM, confirming that bezafibrate has positive effects on mitochondrial phenotype in the BBB *in vivo*.

**Figure 3.**
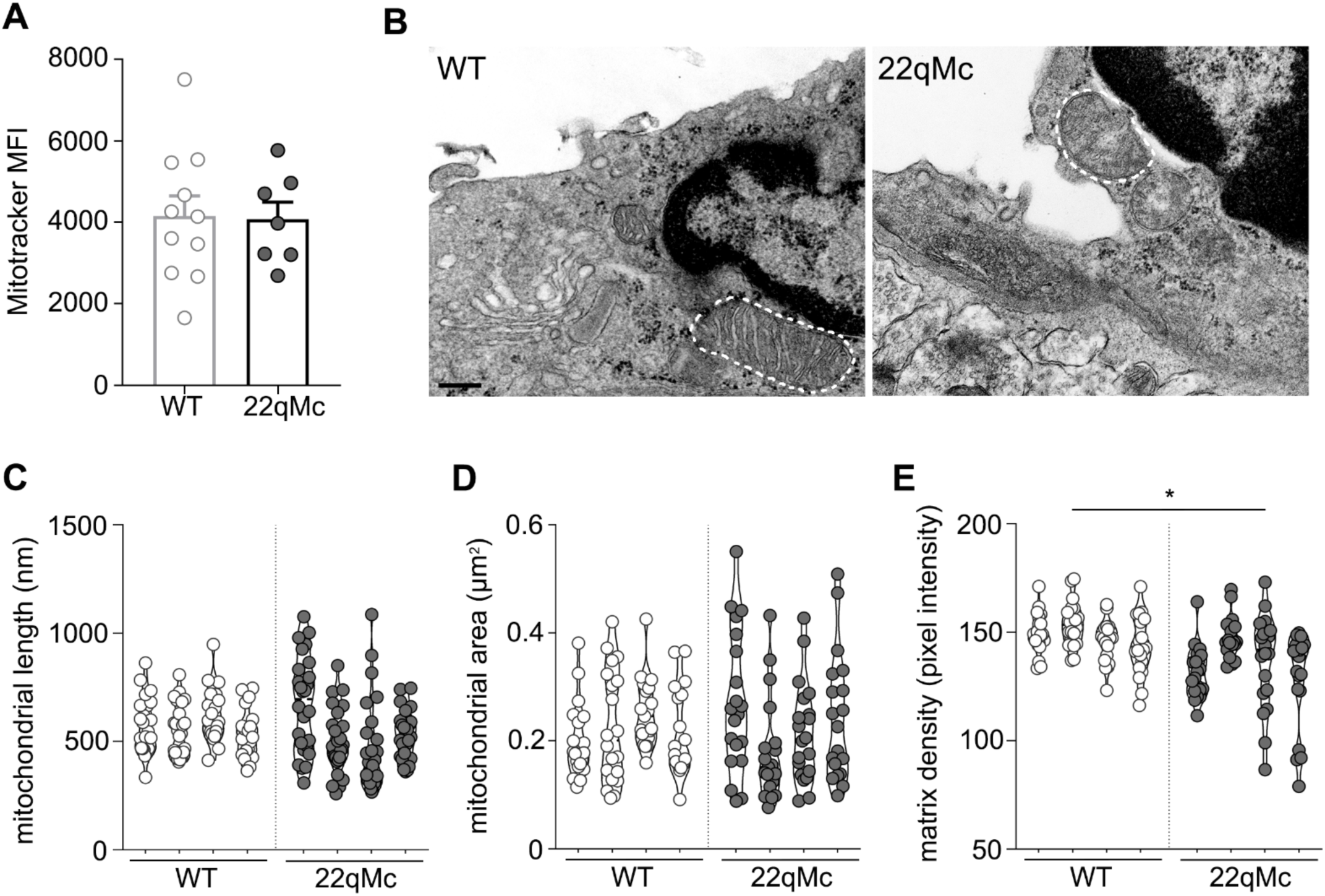
Mitochondria in the 22qMc BBB are dysfunctional. (**A**) Quantification of mitochondrial content by Mitotracker fluorescence (*n* = 7-11 per genotype), and (**B**) Representative images of BBB-EC mitochondria by transmission EM in 22qMc and WT. Quantification of mitochondrial (**C**) length (**D**) area and (**E**) intramitochondrial density in the neurovasculature of 22qMc and WT (*n* = 4, 20 mitochondria per mouse). Error bars ± SEM. *, p < 0.05 by nested t test.

**Figure 4.**
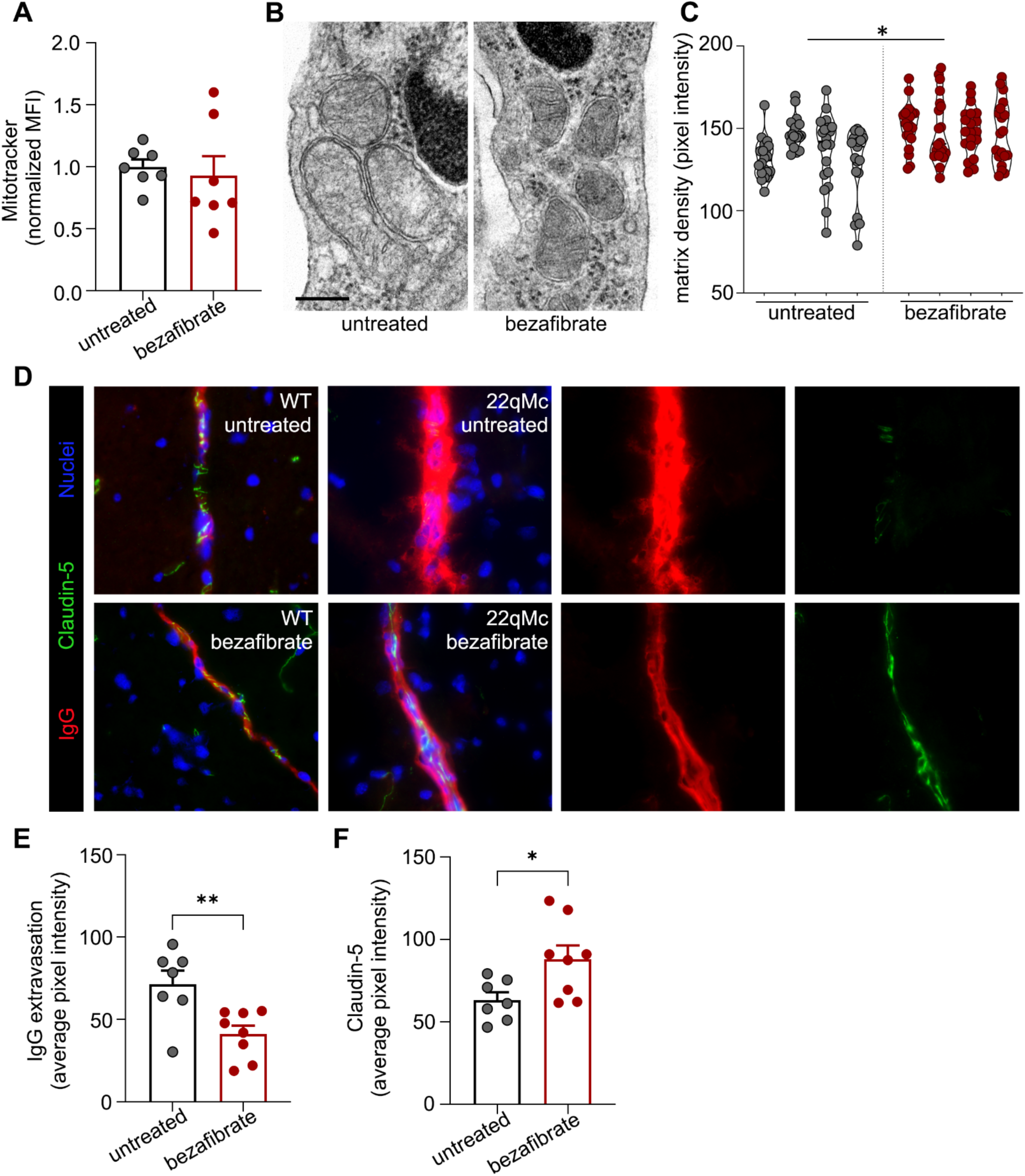
Bezafibrate treatment improves mitochondrial phenotype and barrier function in the 22qMc BBB. (**A**) Quantification of mitochondrial content by Mitotracker fluorescence (*n* = 7). (**B**) Representative EM images and (**C**) quantification of intramitochondrial density in BBB-ECs from treated and untreated 22qMc. (**D**) Representative immunofluorescent staining hippocampal vessels for leakiness (IgG, red) and claudin-5 (green). (**E**) Quantification of BBB leakiness as measured by serum IgG extravasation and (**F**) quantification of claudin-5 in hippocampal vessels in treated and untreated 22qMc (*n* = 7-8). Error bars ± SEM. Scale bar: *, p < 0.05, ** p < 0.01 by nested t test.

We have previously reported increased extravascular serum leakage and decreased claudin-5 expression in the 22qMc BBB, together indicated that barrier function is compromised *in vivo* (*33*). As we found that bezafibrate significantly improved barrier function *in vitro*, we next assessed the effect of bezafibrate treatment on barrier function and claudin-5 expression in the 22qMc BBB. We report that four weeks of bezafibrate treatment significantly decreased extravascular leakage of the serum protein IgG in the hippocampus (Figure 4D-E), indicating improved barrier function in a behaviorally-relevant brain region. We observed a concomitant increase in claudin-5 expression in these same vessels (Figure 4D, F), implicating improved tight junction expression in the effects of bezafibrate on the 22qMc BBB.

To further explore the effects of bezafibrate on the BBB in 22qMc, we performed bulk transcriptomic analysis on BBB-ECs (CD31^+^, CD45^-^, PDGRFb^-^, platelet-derived growth factor receptor b) isolated from untreated 22qMc compared to bezafibrate-treated. We identified 1,051 differentially expressed genes following treatment (Figure 5A). Of these, 335 were significantly upregulated upon bezafibrate treatment, while 375 were significantly downregulated relative to untreated 22qMc (Figure 5B). Within these genes sets, we identified multiple mitochondrial pathways that were impacted by bezafibrate treatment. Specifically, we observed a four-fold increase in *Oip5* transcripts, which encodes Opa-interacting protein 5, a protein that localizes to the mitochondrial outer membrane where it interacts with fusion and fission proteins and regulates mitophagy (*49*). We also observed an increase in genes related to mitochondrial transcription and translation, including *Mrps21* (a mitochondrial ribosomal protein), *Polrmt* (mitochondrial RNA polymerase), and *Mtif3* (mitochondrial translation initiation factor). Interestingly, we observed that the pseudogene for *Mrps21* (*AC166832.2*) was strongly downregulated with treatment.

**Figure 5.**
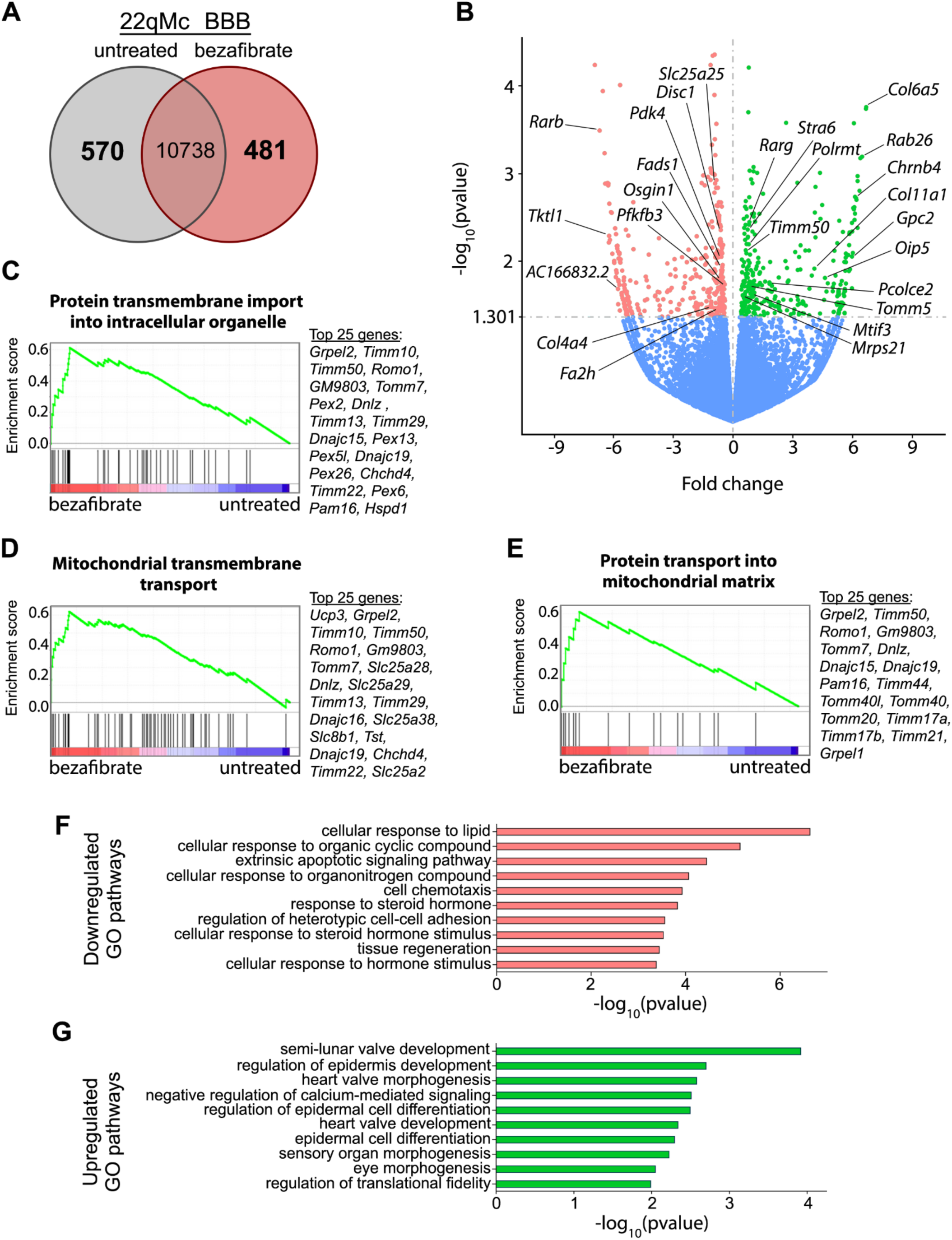
Differential gene expression in the BBB of 22qMc after bezafibrate treatment. (**A**) Venn diagram of differentially expressed genes between untreated and bezafibrate-treated 22qMc. (**B**) Volcano plot of RNA-seq. expression data volcano plot of significantly differentially-expressed genes in flow-sorted CD31^+^CD45^-^ PDGFRb^-^ –BBB-ECs isolated from the brain of 22qMc untreated (*n* = 3) and 22qMc receiving bezafibrate (*n* = 3). Differentially expressed genes with an average fold change of 1051 and P<0.05 were considered significant. The 570 differentially upregulated genes are shown in red and the 481 most downregulated genes denoted in green. Relevant genes are annotated. (**C-E**) Representative gene set enrichment analysis (GSEA) plots of significantly enriched pathways and ranked list of the 20 most-contributing genes (NES = normalized enrichment score). Gene sets with both a P-value < 0.05 and false discovery rate (FDR)<0.05 for GO gene sets were considered significant. Pathway analysis of top (**F**) downregulated and (**G**) upregulated differentially expressed genes using GO enrichment analysis.

We identified multiple genes from diverse metabolic pathways that were downregulated by bezafibrate treatment. We found decreased expression of the gene *Tktl1*, which is known to promote survival during low oxygen conditions, by fermenting glucose into lactate (*50, 51*). Similarly, we observed a decrease in *Pfkfb3* (6-phosphofructo-2-kinase/fructose-2,6-biphosphatase 3), which is involved in stimulating glycolysis (*52*), and *Pdk4* (pyruvate dehydrogenase kinase, isoenzyme 4), which inhibits the oxidation of pyruvate in the mitochondria for use in respiration (*53*). Downregulation of these genes together suggest a decreased reliance on glycolysis for ATP production, or an enhanced reliance on oxygen-dependent respiration. At the same time, decreases in *Fads1* (fatty acid desaturase 1) and *Fa2h* (fatty acid 2 hydroxylase) transcripts further suggest energetic changes in the 22qMc BBB following treatment with bezafibrate. Less strongly downregulated is *Osgin1*, an oxidative stress-induced growth inhibitor (*54*), as was *Disc1*, named for its association with SZ (disrupted in SZ) (*55*). Although the function of *Disc1* is unclear, it is believed to be related to mitophagy, fusion/fission and mitochondrial transport (*56–58*).

We also observed changes in genes promoting vascular function, including high upregulation of pro-angiogenesis gene *Chrnb4*, as well as increased *Ang*, a potent stimulator of angiogenesis (*59–61*). We further found an increase in pro-barrier transcripts, including *Rab26*, which is known to protect cell-cell junctions in other tissues (*62, 63*), *Rarg*, a retinoic acid receptor, as well as the plasma membrane protein *Stra6* (stimulated by retinoic acid), which is responsible for cellular uptake of retinoic acid precursors (*64, 65*). As retinoic acid is known to induce BBB properties (*66, 67*), this is consistent with our findings that BBB integrity is improved by bezafibrate. The most highly upregulated gene following treatment was *Col5a5*, encoding for a component of collagen, while *Col11a1* also strongly upregulated. The extracellular matrix (ECM) supports BBB function (*68, 69*) and its potential role upon bezafibrate treatment is reinforced by the finding of increased *Pcolce2*, procollagen C endopeptidase enhancer 2, which supports maturation of collagen fibrils (*70*), and *Gpc2*, which encodes for a receptor with a high affinity for laminin (*71, 72*), a key component of the ECM at the BBB (*73*).

These observations were supported by unbiased gene set enrichment analysis (GSEA), which reinforced the direct effect of bezafibrate on the mitochondria of the BBB. Specifically, we find that the gene sets enriched in the bezafibrate-treated group are primarily associated with protein transport, most notably “mitochondrial transmembrane transport” and “protein transport into mitochondrial matrix” (Figure 5C-E). Meanwhile, pathway analysis identified GO terms associated with vascular development and morphogenesis as upregulated in 22qMc treated with bezafibrate, including semi-lunar valve development, pulmonary valve development and morphogenesis, and heart valve development and morphogenesis (Figure 5G. Together, our transcriptomics results indicate that bezafibrate treatment impacts the mitochondria in the BBB of 22qMc and is associated with acquisition of a pro-vascular phenotype promoting mitochondrial and barrier function.

### Bezafibrate treatment improves social memory behavior in 22qMc

We next aimed to assess the behavioral impacts of bezafibrate treatment and the associated improvement in BBB integrity. As we found that bezafibrate enhanced BBB function in the hippocampus in 22qMc, we chose to investigate behavioral performance in social memory, a hippocampal-dependent task (*74*). Performance in this task is known to be affected by hippocampal BBB function, as induced claudin-5 knockdown within the hippocampus is sufficient to induce deficits in social memory (*10*). Social memory is a reported behavioral deficit in 22qMc (*75, 76*), and we have found that the BBB is impaired in 22qMc (*33*). Therefore, we hypothesized that BBB deficits contribute to social memory impairment in 22qMc. To determine whether improving BBB integrity is associated with improved social memory, we treated 22qMc with 0.5% bezafibrate-enhanced chow for 4 weeks prior to behavior assessment (Figure 6A). As has been previously published, we found that 22qMc perform at control levels in the social preference task but do exhibit impairment in social memory (Figure 6B-C) (*75, 76*). The deficit in social memory was restored in 22qMc treated with bezafibrate, as they show a significant increase in the percentage of time spent interacting with the novel mouse, compared to untreated 22qMc (Figure 6C). Indeed, bezafibrate-treated 22qMc perform indistinguishably from WT mice.

**Figure 6.**
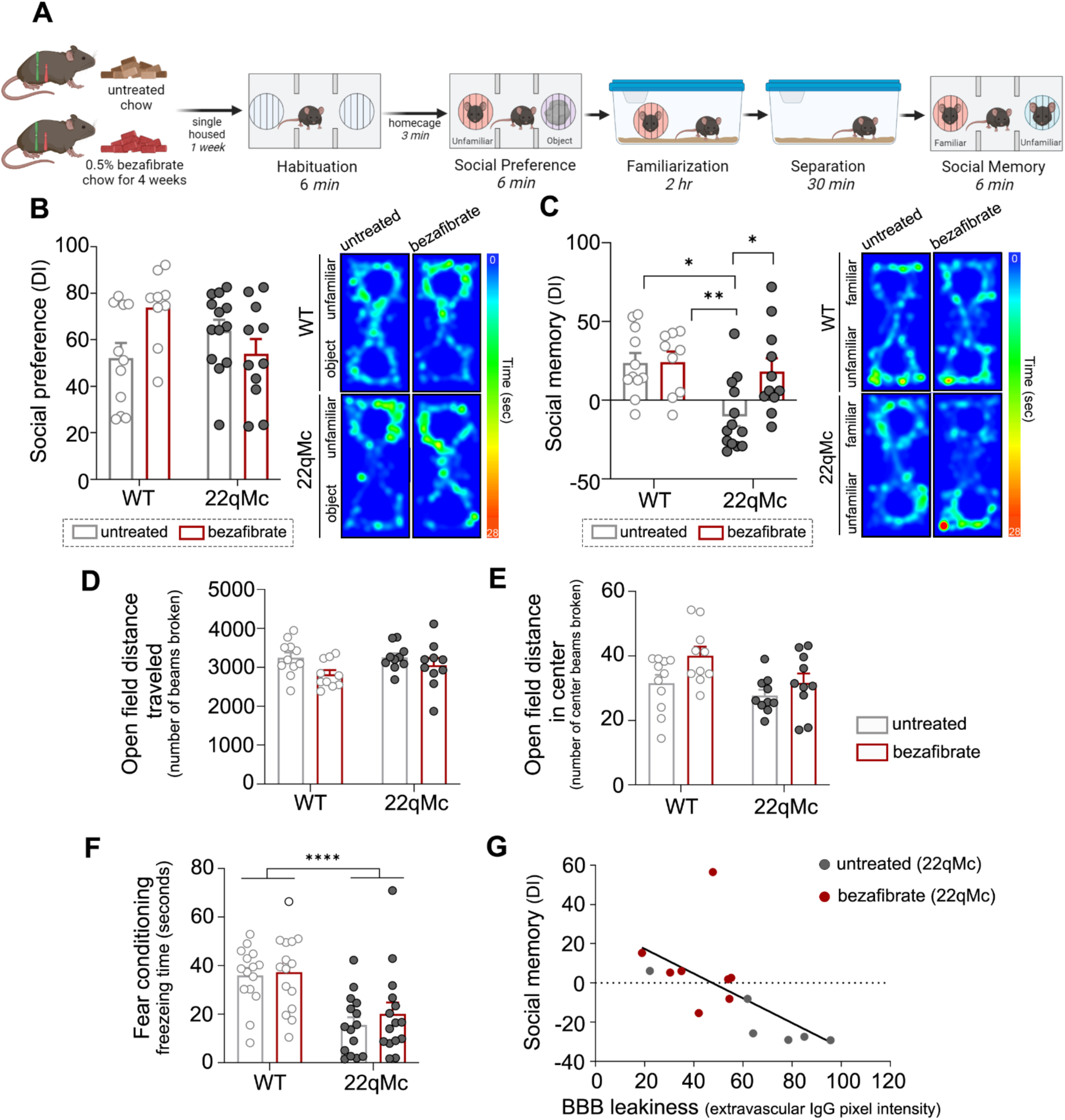
Bezafibrate treatment improves social memory behavior in 22qMc. (**A**) Schematic depicting the protocol to carry out the social preference and social memory behavioral tasks. (**B**) Social preference and (**C**) social memory behavioral task performance (left) and representative heatmaps of mouse activity during the task (right) (*n* = 11-13 per group). (**D**) Hyperactivity as measured by total number of beams broken during the open field test. (**E**) Anxiety-like behavior as measured by number of beams broken in the center of the open field. (**F**) Fear conditioning as measured by freezing time during the recall phase of the task (not shown, no difference in freezing time prior to the task, no difference in freezing time during the acquisition phase of the task) (*n* = 15 per group). (**G**) Correlation of social memory performance and BBB leakiness in 22qMc (R = -0.6275, p < 0.05 by Pearson’s correlation). Error bars ± SEM. *, p < 0.05, ** p < 0.01, **** p < 0.0001 by two-way ANOVA unless otherwise indicated.

To confirm that 22qMc do not exhibit hyperactivity, or that bezafibrate does not impact an anxiety-like phenotype that could account for these results, we also assessed treated and untreated 22qMc and WT in open field. We report no difference in total distance traveled or center arena activity, demonstrating no effect of genotype or treatment on these behaviors (Figure 6D-E). We also tested the specificity of the behavioral rescue by assessing contextual fear conditioning, another hippocampal-dependent task that is known to be affected in 22qMc but is not associated with alterations in BBB integrity (*77*). As expected, we found that bezafibrate treatment had no effect on fear conditioning in 22qMc, as both treated and untreated mice failed to recognize the chamber in which they had previously received a footshock (Figure 6F). Together these data suggest a specific effect of bezafibrate on social memory, a task previously associated with BBB integrity in the hippocampus.

Based on the published link between hippocampal BBB integrity and social memory performance (*10*), we hypothesized that BBB function was associated with the ability to recognize the novel mouse. To address this, we compared the extent of BBB leakiness in the hippocampus (as measured by the fluorescent intensity of extravascular IgG) to the social discrimination score in the memory phase of the behavioral task (as measured by the relative percentage of time spent interacting with the novel mouse). We found a significant negative correlation between BBB leakiness and social memory performance, such that stronger IgG extravasation was associated with worse social memory (Figure 6G).

## Discussion

Mitochondrial deficits are closely tied to SZ generally and 22qDS specifically (*17, 37, 78*). While much work has focused on the contributions of specific genes in the 22q11.2 deletion, most notably *Mrpl40* and *Txnrd2* (*14, 32, 38*), here we focus on how the deletion as a whole contributes to a translationally important and therapeutically-targetable deficit in mitochondrial function and BBB integrity. Notably, BBB-ECs harbors a higher mitochondrial content compared to peripheral endothelial cells (*20*), suggesting specialized metabolic requirements of the BBB. Indeed, others have identified a link between mitochondrial insults and barrier integrity, primarily in the context of stroke (*22, 23, 79, 80*). Yet, relatively little is known about mitochondrial function in the BBB, and mechanistic associations between mitochondrial respiration and BBB integrity have not yet been explored. For this reason, we chose to study the relationship between mitochondrial function and barrier integrity in the context of 22qDS, a translationally relevant and uniquely appropriate model to address such questions.

Here, we report mitochondrial dysfunction in the 22qDS BBB, a model of genetic susceptibility to SZ that we have previously reported exhibits barrier impairment (*33*). We probed mitochondrial function in 22qDS iBBBs, and report decreased respiration by both the Resipher and Seahorse assays. We identified a deficit in ATP production *in vitro*, suggesting that mitochondrial dysfunction can impact the cellular energetics of the BBB in 22qDS. As technical approaches to measure mitochondrial respiration *in vivo* are lacking, we were not able to identify deficits in mitochondrial respiration in the 22qMc BBB. Instead, we investigated mitochondrial structure by electron microscopy, and identified that the intramitochondrial density is reduced in the 22qMc BBB compared to WT. While experimental evidence linking such changes in the mitochondrial phenotype by EM to discrete aspects of mitochondrial biology (such as mitochondrial membrane potential, calcium flux, density, and function of electron transport chain complexes, for example) is lacking, together our findings suggest that mitochondria are dysfunctional in the 22qDS BBB.

To mechanistically address the relationship between barrier function and mitochondrial dysfunction, we treated with the PPAR agonist bezafibrate to increase mitochondrial respiration. We chose this compound for a number of reasons: (1) treatment with PPAR agonists have been shown to improve BBB integrity *in vitro* following oxygen-glucose deprivation and *in vivo* following ischemic stroke (*80, 81*), indicating that it is capable of impacting BBB function, (2) PGC1α, the PPAR downstream signaling molecule has been reported to increase capacity for ATP production in BBB-ECs (*82*), indicating its potential to improve respiration in the BBB, (3) bezafibrate treatment significantly improved ATP production in the context of 22qDS (*39*), suggesting that treatment can overcome the mitochondrial dysfunction conferred by the deletion, (4) fibrates are believed to be inefficient at crossing the BBB (*79*), an additional advantage given our goal of studying biological and behavioral effects on the BBB, (5) PPAR agonists have shown positive outcomes in preclinical studies of BBB-associated neurological diseases, including stroke, epilepsy, SZ and Alzheimer’s disease (*83–87*), reinforcing the broad translational applications of our findings and (6) bezafibrate is an FDA-approved compound currently being used for treatment of metabolic disorders, reflecting the translational goals of this study.

Using the Seahorse mitochondrial stress test, we demonstrate that bezafibrate treatment enhances mitochondrial function in the 22qDS BBB. While the mechanisms behind bezafibrate’s impact on mitochondrial function in the 22qDS BBB need to be investigated further, our data suggest that bezafibrate acts to improve mitochondrial respiration via changes in mitochondrial dynamics. It has been proposed that mitochondrial fusion and fission may be a mechanism by which cells can regulate the health of the existing pool of mitochondria in order to enhance the respiratory capacity (*88*). Indeed, we find that bezafibrate improves mitochondrial phenotype in the 22qMc BBB, as evidenced by restoration of the mitochondrial matrix density by EM. These findings were reinforced by our sequencing results, which identified multiple upregulated transcripts that have been linked to mitochondrial dynamics. This includes *Oip5,* as well as *Pdf4*, whose overexpression has been found to promote mitochondrial fission (*89*), and *Disc1*, which was shown to interact with and regulate the prominent fission protein, Drp1 (*56, 57*). We also found a significant effect of bezafibrate on transcripts associated with protein transport within intracellular organelles, specifically the mitochondria. This was accompanied by increases in transcripts related to mitochondrial protein synthesis, which would be required because of changes in mitochondrial dynamics and could be associated with the improved mitochondrial health observed following treatment. While transcripts more directly associated with fusion and fission (such as those studied in Figure 2D) did not emerge in our sequencing analysis, this is likely due to the dramatic differences in time course of treatment *in vitro* compared to *in vivo*. Nevertheless, our findings support the hypothesis that bezafibrate improves mitochondrial respiration by affecting mitochondrial dynamics and together our findings suggest that bezafibrate improves mitochondrial function in the 22qDS BBB.

We identified that bezafibrate’s effects on mitochondrial respiration coincided with a significant improvement in barrier function and tight junction expression in the 22qDS iBBB, indicating that mitochondrial function directly affects barrier integrity *in vitro*. Importantly, we cannot make this assumption *in vivo*, as bezafibrate is known to impact metabolism and inflammation, with effects on multiple other organ systems including the liver. However, our transcriptomic analysis point towards a direct effect of bezafibrate at the level of the BBB, as we observe upregulation of mitochondrial gene sets in BBB-ECs isolated from bezafibrate-treated 22qMc. Thus, our findings implicate bezafibrate treatment directly in the pro-barrier effects on the BBB. The mechanistic interactions linking mitochondrial respiration and ATP production to barrier integrity and claudin-5 expression remain unclear, and these aspects should be addressed in future studies. Together, our results identify the mitochondria as a novel therapeutic target by which barrier function can be enhanced in 22qDS, and we speculate that mitochondrial deficits contribute to the alterations in BBB integrity that we have previously reported in this condition.

Here we focused on the relationship between BBB integrity and hippocampal-dependent tasks. Structural changes in the hippocampus are observed in neuropsychiatric diseases, and patients exhibit impairment in hippocampal-dependent tasks, making this is a translationally-relevant brain region (*90–92*). For this reason, we assessed social memory performance, a task known to be dependent on hippocampal functioning (*74*). Prior reports indicate that social memory, but not social preference, is directly modulated by BBB integrity in the hippocampus (*10*). As 22qMc exhibit BBB impairment localized to the hippocampus (*33*), and recapitulate this behavioral phenotype (*75*), we hypothesized that BBB dysfunction contributes to social memory deficits in 22qMc. Given that we observed enhanced BBB integrity in the hippocampus following bezafibrate treatment, we anticipated that bezafibrate would improve a BBB-dependent task in 22qMc. Indeed, we found that bezafibrate treatment improved social memory performance. However, it is important to note that we cannot confirm that effects of bezafibrate on BBB function alone was sufficient to alter behavior. To our knowledge, there are no pharmaceutical interventions that specifically target mitochondrial function in the BBB, limiting our ability to mechanistically address this question. In the context of our findings that BBB integrity is significantly correlated with social memory and the published literature on these topics, our data strongly suggest a role for hippocampal BBB integrity in contributing to social memory in 22qDS.

For the first time, we identify the mitochondria as a therapeutic target to modulate barrier function of the BBB in neuropsychiatric disease. Here, we report mitochondrial dysfunction in the 22qDS BBB characterized by decreased mitochondrial respiration, impaired ATP production and enhanced ROS accumulation. We identify bezafibrate as a treatment that improves mitochondrial health and function in the 22qDS BBB, potentially via its effects on altering mitochondrial dynamics. We found that these beneficial effects on the mitochondrial are accompanied by enhanced barrier function of the 22qDS BBB both *in vitro* and *in vivo*. Finally, we link improved BBB function to rescue of neuropsychiatric behavior, as we report that treatment with bezafibrate restores social memory in the murine model of 22qDS. Together, our findings posit mitochondrial dysfunction as a contributor to BBB impairment in neurological disease and identify a novel therapeutic mechanism by which the BBB may be targeted in these conditions.

## Materials & Methods

### Human iPSC**-**derived iBBB cultures

All human experiments were conducted under Internal Review Board approval by the University of Pennsylvania and the Children’s Hospital of Philadelphia (CHOP). Human iPSC lines derived from 4 patients with 22qDS and SZ and 4 age/sex matched healthy controls were generously received from Dr. Sergiu P. Pașca, Stanford University, Stanford, California. Human iPSCs were differentiated into iBBBs, stored, and cultured following the protocols previously published (*33, 93, 94*). For experiments, plates were coated with a collagen/fibronectin mixture containing: 50% H_2_O, 40% 1mg/mL collagen from human placenta (Sigma) and 10% fibronectin from bovine plasma (Sigma). Cells were counted and plated according to Table 1. Cells were cultured in EC medium (human endothelial serum-free media (Thermo) containing B-27 Supplement without antioxidants (Thermo) as previously reported (*33, 94*). Bezafibrate (Thermo) was diluted in 0.02% DMSO (100 μM), and treatment was initiated 24 hours after plating. Outcomes were measured 36-48 hours later unless otherwise indicated. All experiments were conducted in a paired format, consisting of age- and sex-matched 22qDS/HC pairings, or treated/untreated conditions of 22qDS.

**Table 1.** iBBB culturing specifications.

| **Experiment** | **Cell Density** | **Media Volume** | **Plating Substrate** |
| --- | --- | --- | --- |
| TEER | $1.75 \times 10^5$ | 400 μL | 8W10E+ electrode arrays<br>(Applied Biophysics, Troy, NY, USA) |
| Immunofluorescence | $8 \times 10^4$ | 300 μL | 8 well chamber slide (ibidi) |
| Flow cytometry<br>qPCR | $3 \times 10^5$ | 1 mL | Nunclon™ Delta Multidishes, 12 well (ThermoFisher Scientific) |
| Resipher | $6 \times 10^4$ | 100 μL | Nunclon™ Delta Multidishes, 96 well flat bottom (ThermoFisher Scientific) |
| Seahorse | $3 \times 10^4$ | 100 μL | Seahorse XF96 V3 PS cell culture microplates (Agilent) |

### iBBB Functional Metabolic Assays

Oxygen consumption in 22qDS and matched HC iBBBs were measured using Resipher (Lucid Scientific) over the course of 4 days. Data represents the average of 4-8 technical replicates for each iBBB line. Seahorse mitochondrial stress test was performed using the Agilent Bioanalyzer with inhibitors at the following concentrations: oligomycin (1.5 μM), FCCP (1 μM), rotenone (1 μM), antimycin A (1 μM). Data was analyzed using Seahorse Wave Desktop Software (Agilent). Data for each phase of the trial was compiled from 3-4 time points per phase and represents the average of 4-8 technical replicates per iBBB line. Unpaired t test was used to assess differences between control and 22qDS lines at each time point in Resipher and for each phase of Seahorse; paired t test was used to analyze treated and untreated 22qDS lines for each phase of Seahorse.

### iBBB TEER

The electrical properties of confluent iBBB monolayers were measured as previously described (*33, 95*). Electric Cell-substrate Impedance Sensing (ECIS) methodology was employed making using of the ECIS Zϴ instrument and 8W10E+ electrode arrays (Applied Biophysics, Troy, NY, USA), and resistance was measured at 4000 Hz. 100 μM bezafibrate was added to the media 2-3 hours prior to confluency (measured as the maximum resistance), and maximal TEER effect was assessed 36 hours later. Each experiment consisted of 3-5 technical replicates, and each 22qDS line was repeated in two separate experiments. Average fold change of the bezafibrate-treated TEER curve was normalized to the average fold change of the untreated TEER curve and was analyzed using two-way ANOVA. Treated and untreated 22qDS lines (n = 4) were analyzed by paired t test.

### qPCR

RNA was isolated using RNeasy kit (Qiagen) for analysis of relative genetic expression, DNA was isolated using DNeasy Blood and Tissue kit (Qiagen) for analysis of mitochondrial DNA content. Samples were run in triplicate and comparative CT was measured on QuantStudio 3 Real Time PCR System (Thermo) using Syber Green master mix. Primer sequences are listed in Table 2. For quantification of mitochondrial DNA, each of 4 mitochondrial genes was normalized to each of 2 nuclear genes; the average of these values is presented as the mitochondrial DNA copy number. All samples assessing RNA expression were normalized to GAPDH expression and fold change relative to the average of the untreated samples was calculated.

**Table 2.**
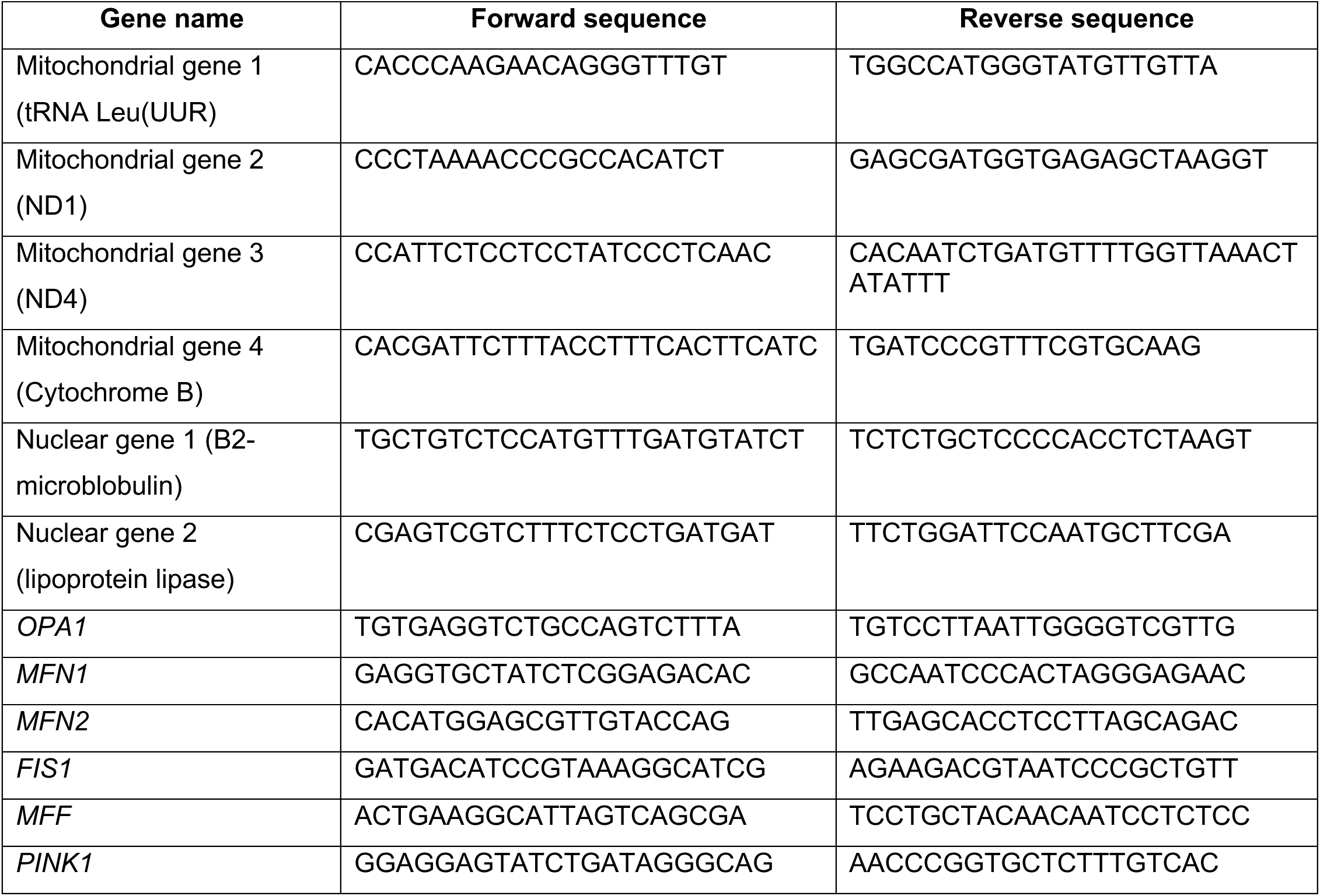
Primer sequences for qPCR analysis.

### iBBB immunofluorescence

Cells were fixed with 70% ethanol, permeabilized with 1X TBS + 0.025% Tween, blocked with 10% normal donkey serum and subsequently stained with rabbit polyclonal antibodies against claudin-5 (Life Technologies) diluted 1:200 in 3% normal donkey serum overnight at 4°C. Secondary antibody Alexa Fluor® 488 AffiniPure Fab Fragment Donkey Anti-Rabbit IgG (Jackson Immunoresearch) was diluted 1:300 and incubated in 3% normal donkey serum before mounting with Gelvatol containing Hoescht (Invitrogen). Cells were imaged using a Leica SP5 confocal microscope (Leica Microsystems). Images were obtained using an Olympus IX83 set up for brightfield and fluorescence, and equipped with a motorized X, Y, Z stage and a spinning disk confocal head (X-Light V2, Crestoptics s.r.l., Rome, Italy) using a Hamamatsu R2 cooled CMOS camera (Hamamatsu City, Japan) operated by the MetaMorph software (Molecular Devices, LLC, Sunnyvale, CA).

### Mitochondrial dyes

To quantify cellular ROS in mouse BBB-ECs and in human iPSC-derived iBBBs, cells were stained with 1 μM CellRox (Thermo) for 1 hr at 37°C, while mitochondrial ROS was quantified by staining with 1 μM MitoSox (Thermo) for 15 min at 37°C. Mitochondrial content was measured using Mitotracker by staining with 1 μM for 30 min at 37°C. BBB-ECs were subsequently stained for endothelial selection markers, and samples were run on an LSR Fortessa and mean fluorescent intensity was analyzed using FlowJo 10 (BD Biosciences).

### Mice

All animals were cared for in accordance with the guidelines of the National Institutes of Health and procedures were approved by the University of Pennsylvania Institutional Animal Care and Use Committees. 12–16-week-old male and female Df(h22q11)/+ and wild type (WT) littermates (Taconic) were maintained on a reverse 12 hr light/dark cycle with ad libitum chow and water. In bezafibrate treatment experiments, mice were given 0.5% bezafibrate enriched chow (TestDiet) or irradiated base chow (Animal Specialties and Procurement) for 4 weeks ad libitum. Treatment time point was based on other experiments reporting CNS outcomes of orally administered bezafibrate. Mice were anesthetized by CO_2_ and transcardially perfused with 1X PBS. Brains were either frozen for whole brain sectioning/western blotting/qPCR or kept in RPMI (1X) + GlutaMAX (Gibco) + 1X MEM NAA for isolation of BBB-ECs. Data was initially analyzed separately and upon confirmation that there was no effect of sex, data from males and females were combined.

### Social Behavior

Social preference and social recall were performed as previously described (*75*). In brief, mice were habituated to the three-chamber arena with empty cylinders for 6 minutes, then returned to the home cage for 3 minutes. The social preference phase followed where the experimental mouse explored a novel juvenile mouse in one cylinder and an inanimate object in the other cylinder for 6 minutes. Mice were returned to the home cage to reinforce familiarize with the same juvenile for 2 hours. The mice were separated for 30 minutes prior to the recall phase. During the recall phase, the experimental mouse is returned to the arena with the now familiar mouse in one cylinder and a novel juvenile mouse in the other chamber. Cues were counterbalanced across experiments. Interaction time with stimuli was measured using ANYMaze video tracking software, and discrimination scores were calculated: 100 x (novel mouse exploration - familiar mouse or object exploration) / total exploration time). Mice that climbed on the cylinders were excluded from analysis. Mice performing more than two standard deviations from the group mean (*n* = 1 mouse in each untreated WT and bezafibrate-treated WT groups) were excluded from analysis. Two mice were excluded for not registering any interaction time with one of the stimuli during one of the trial phases.

### Open Field

Spontaneous activity in an open field arena was used to assess ambulation and rearing as part of a general health assessment. Mice were habituated to the procedure room for 30 minutes before the trial. To start the trial, a mouse was placed in the center of a Plexiglas arena (14 inches x 14 inches) with 4 clear 18-inch walls, fitted with infrared emitters and detectors. Activity was collected as beam breaks using a Photobeam Activity System (San Diego Instruments). Horizontal, rearing, center and peripheral beam breaks were collected in a 10-minute trial.

### Contextual Fear Conditioning

Contextual conditioning is a hippocampus-dependent form of learning in which an aversive stimulus (foot shock) becomes associated with a unique environment, the conditioning chamber. Upon subsequent exposure to the conditioning chamber, mice that associate the context with the aversive stimulus will freeze in response; reduced freezing suggests a memory impairment. Mice were handled for 2 min on three consecutive days prior to the acquisition trial. For acquisition, mice were placed in a conditioning chamber (Med Associate) within a sound-attenuating cabinet. A 1.25 mA foot shock was delivered between 148–150 seconds of a 180-second trial. Twenty-four hours after acquisition, mice were returned to the conditioning chamber for a 300-second recall trial to assess long-term memory. High speed, digital recordings of all trial were processed by automated analysis with VideoFreeze software. The percent time freezing was defined as the mouse being motionless except for respiratory movements.

### Immunofluorescent staining

Sagittal brains were cryosectioned at 8 μm, mounted onto charged slides and immunostained as previously described. In brief, brains were fixed in acetone and 70% ethanol, permeabilized with 1X TBS + 0.025% Tween and blocked with 10% normal donkey serum. Primary antibody rabbit polyclonal antibodies against claudin-5 (Life Technologies) was diluted 1:300 in 3% normal donkey serum and incubated overnight at 4°C. Sections were washed with 1X TBS and incubated with secondary antibodies Alexa Fluor® 488 AffiniPure Fab Fragment Donkey Anti-Rabbit IgG (Jackson Immunoresearch) and Rhodamine Red^TM^-X AffiniPure Fab Fragment Donkey Anti-Mouse IgG (Jackson Immunoresearch) in 3% normal donkey serum at room temperature. Nuclei were permeabilized with 1X TBS + Triton X-100 and slides were mounted with Gelvetol containing Hoescht nuclear dye. Sections were imaged on Leica widefield microscope (Leica Microsystems) and all immunofluorescent analyses were performed blinded and using ImageJ (NIH), as previously described (*33*).

### Electron Microscopy

Samples were collected as stated above and processed for electron microscopy (EM) as previously published (*96*). brain samples (∼1 mm^2^) were cut in 90-nm sections, collected on a 150-mesh grid and stained with saturated aqueous uranyl acetate and Reynolds lead citrate. Transmission EM was carried out at the electron microscopy core of the University of Texas Health Science Center of San Antonio (UTHSCSA) using a JEOL 1400 electron microscope (JEOL, Peabody, MA). Image analysis was performed blinded using ImageJ (NIH).

### Isolation of BBB-ECs for flow cytometry

Brains were dounce homogenized in RPMI (1X) + 10% Cosmic Calf Serum (Hyclone) and digested with 100 mg/mL collagenase/dispase (Roche) and 1 mg/mL DNase I (Roche) for 25 min at 37°C with agitation. BBB-ECs were isolated through density centrifugation with 22% Percoll (Sigma). Cells underwent mitochondrial dye loading as described above and were subsequently stained with fluorescently-tagged endothelial positive and negative selection markers for flow cytometry.

### Isolation of BBB-ECs for RNA sequencing

Brains ECs were isolated using a protocol adapted from Munji et al., 2019 for sequencing experiments (*97*). For RNA sequencing, 3 brains of the same sex, genotype and treatment conditions were pooled into each sample (n = 3 samples per treatment condition). Brains were minced in PBS and subsequently digested in papain (Worthington) at 35°C for 90 minutes. Brains were triturated in trypsin inhibitor (ovomucoid; Worthington) and the cell suspension was digested in collagenase (Worthington) and neutral protease (Worthington) for 30 minutes at 35°C. Myelin was removed through density centrifugation with 22% Percoll. Cells were stained with FITC anti-mouse CD45 (Biolegend, 1:50), Zombie Violet (Biolegend 1:500), PE anti-mouse PDGFRb (Biolegend, 1:100), and APC-Cy7 anti-mouse CD31 (Biolegend, 1:50) for 30 minutes at 4°C in PBS. Live endothelial cells (CD31^+^, CD45^-^, PDGFRb^-^) were isolated by sorting on an Influx Cell Sorter (BD Biosciences).

### RNA sequencing

Cells were sorted into RNA lysis buffer (RLT buffer, Qiagen) and total RNA was isolated using the RNeasy Plus Micro Kit (Qiagen) and sample quality was assessed on 4200 TapeStation (Agilent) as previously performed (*98*). mRNA library was prepared using the SMART-Seq HT kit (Clontech). Illumina indexes were added to cDNA using the Nextera XT DNA library preparation kit (Illumina). RNA-seq libraries were subjected to single end 75 bp read sequencing on an Illumina NextSeq 500 sequencer by Novogene (Beijing, China). Analysis was performed as previously described (*98*). In brief, processed reads were de-duplicated and mapped to the mouse transcriptome using Salmon (*99*) with default settings and using the Ensemble mouse GRCm38 cDNA annotation as a reference transcriptome. Transcripts were filtered by discarding any with TPMs <1. Transcripts were then converted to genes, with the transcript with the highest average TPM being used. Differentially expressed genes were defined as those with both a >1.5 average fold change and a *t*-test *P*<0.05 when comparing control BBB-ECs to bezafibrate BBB-ECs samples.

### Statistics

Unpaired t test was used for iBBB experiments in which controls were compared with 22qDS, as well as all analyses of mouse groups by immunofluorescent staining and flow cytometry. Paired t test was used for iBBB experiments in which differences between treated and untreated 22qDS were assessed. Two-way ANOVA was used to analyze representative Resipher time course and representative normalized TEER time course. Two-way ANOVA was also used to determine significance in all behavioral analyses, and Grubb’s (alpha < 0.05) was used to exclude outliers. Nested t test was used to analyze the EM data. Correlation significance was assessed using Pearson’s correlation. All data was analyzed using GraphPad Prism 9. Only relevant comparisons are shown, values are presented as mean ± SEM unless otherwise indicated. P values are * *P* ≤ 0.05, ** *P* ≤ 0.01, *** *P* ≤ 0.001, **** *P* ≤ 0.0001.

## Abbreviations

(22qDS): 22q11.2 deletion syndrome
(22qMc): 22q11.2 deletion syndrome mouse model
(ATP): Adenosine triphosphate
(BBB): Blood-brain barrier
(CCCP): Carbonyl cyanide m-chlorophenyl hydrazone
(CNS): Central nervous system
(ECs): Endothelial cells
(ECAR): Extracellular acidification rate
(ECM): Extracellular matrix
(HC): Healthy control
(iPSC): Induced pluripotent stem cell
(iBBB): Induced pluripotent stem cell-derived blood-brain barrier
(MFF): Mitochondrial Fusion Factor
(MFN): Mitofusin
(OCR): Oxygen consumption rate
(OPA1): Optic atrophy gene 1
(PDGFRb): Platelet derived growth factor receptor b
(PPAR): Peroxisome proliferator activated receptor
(PGC1α): Peroxisome proliferator-activated receptor gamma coactivator 1 α
(ROS): Reactive oxygen species
(SZ): Schizophrenia
(TEER): Transendothelial electrical resistance
(WT): Wild type

## Acknowledgements

We would like to thank Tim O’Brien and the Neurobehavioral Core for their maintenance of the 22qMc colony, the use of their facilities and equipment, as well as sharing their expertise in behavioral implementation and analysis. Special thanks to Dr. Meagan McManus for critical feedback on the content of this manuscript. We thank Dr. Amit Bar-Or for allowing us the use of the LSR Fortessa, as well as the Penn Cytomics and Cell Sorting Core for machine upkeep and their sorting services. Special thanks to Dr. Barbara Hunter (University of Texas Health Science Center at San Antonio) for electron microscopy support and guidance. We appreciate the advice obtained from Drs. Charles Vite and Gary Swain (Clinical Sciences and Advance Medicine - Penn Vet) on confocal microscopy and Gordon Ruthel at the Penn Vet Imaging Core of the University of Pennsylvania for assistance on scoping and image analysis. Thank you to the Zhang lab at Emory University and the Daneman lab at UC San Diego for freely sharing their BBB isolation protocols. We are indebted to the Kahn lab at UPenn for the use of their ECIS instrument and to the Henao-Mejia lab at UPenn for sharing access to the Resipher device. We appreciate Ms. Renee Rawls and the animal care technicians and veterinarians for support with the murine experiments. We appreciate the support of the Beiting lab and the Center for Host Microbe Interactions for their help and advice in optimizing and assessing RNA quality for sequencing experiments. Finally, we thank the Islet Cell Biology Core for support and implementation of the Seahorse Bioanalyzer experiments.

