## Supplementary data for "Neurovascular mitochondrial susceptibility impacts blood-brain barrier function and behavior"

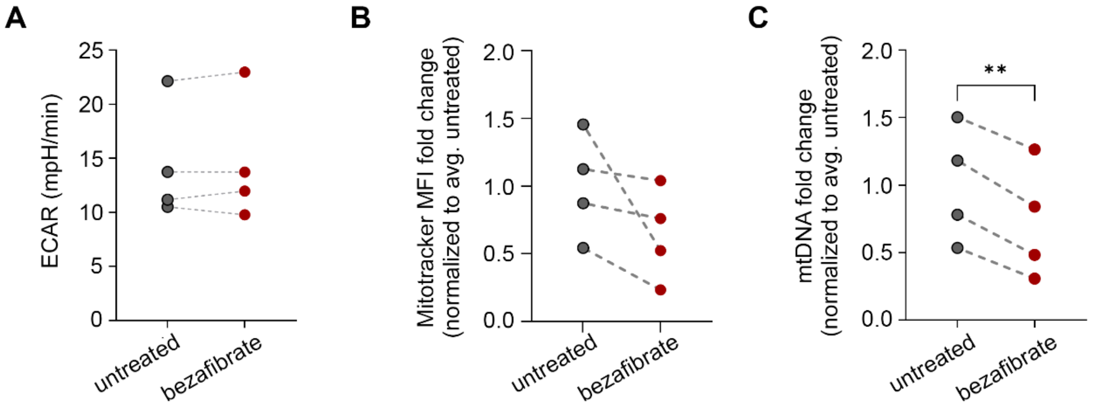


**Supplementary Figure 1. Bezafibrate does not enhance glycolysis or change mitochondrial mass** (A) ECAR measured by Seahorse XFe96 Analyzer to assess glycolysis (*n* = 4 22qDS lines, 7 replicates per condition, not significant by paired t test). (B) Quantification of mitochondrial DNA copy number and (C) Mitotracker mean fluorescent intensity (*n* = 4 22qDS lines, paired t test). Error bars ± SEM. ** p < 0.01

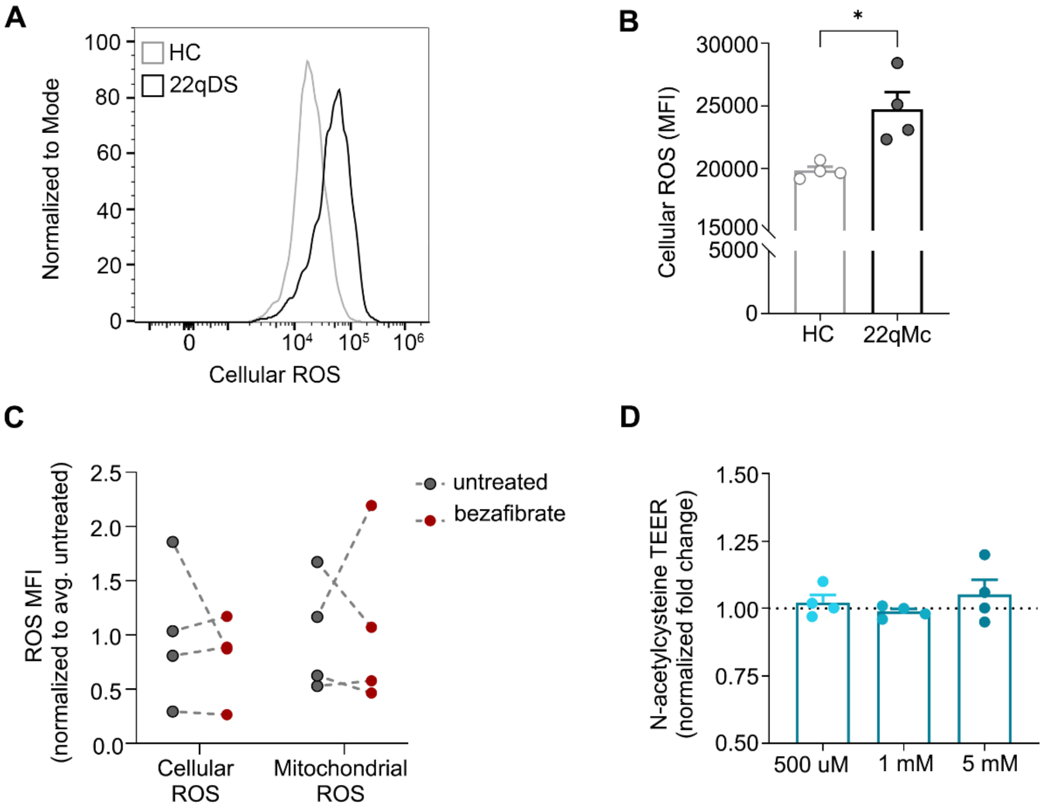


**Supplementary Figure 2. Enhanced ROS accumulation does not drive barrier integrity in the 22qDS iBBB.** (A) Representative histogram of CellRox staining in 22qDS and HC iBBBs by flow cytometry. (B) Cellular ROS quantified as mean fluorescent intensity (*n* = 4 lines per genotype, unpaired t test). (C) Fold change of mean fluorescent intensity of cellular and mitochondrial ROS following treatment with bezafibrate, normalized to the untreated condition (*n* = 4 22qDS lines, not significant by paired t test). (D) Quantification of the TEER fold change at 36 hours following treatment with antioxidant N-acetyl cysteine, normalized to the untreated (*n* = 4 22qDS lines, 3-4 replicates per condition, not significant by Wilcoxon matched pairs signed rank test). Error bars ± SEM. * $p < 0.05$.
